# Distinct ossification trade-offs illuminate the shoulder girdle reconfiguration at the water-to-land transition

**DOI:** 10.1101/2023.07.17.547998

**Authors:** Janet Wei, Thomas W.P. Wood, Kathleen Flaherty, Alyssa Enny, Ali Andrescavage, Danielle Brazer, Dina Navon, Thomas A. Stewart, Hannah Cohen, Anusha Shanabag, Shunya Kuroda, Ingo Braasch, Tetsuya Nakamura

**Author notes:** Address correspondence to: Tetsuya Nakamura, Department of Genetics, Rutgers the State University of New Jersey, Piscataway, LSB 224, 145 Bevier Rd, Piscataway, NJ, 08854. these authors equally contributed to this paper (Janet Wei, Thomas W.P. Wood, Kathleen Flaherty).

## Abstract

The mechanisms of the pectoral girdle transformation at the origin of terrestrial locomotion in vertebrates remains an outstanding problem in evolutionary biology^1^. The loss of dermal bones and the enlargement of endochondral bones resulted in the disarticulation of the pectoral girdle from the skull and the formation of the neck during the fish-to-tetrapod transition^2–5^. Despite the functional implications of this skeletal shift in the emergence of terrestrial vertebrates, the underlying genetic-developmental alterations have remained enigmatic. Here, we discovered that in zebrafish pectoral girdle mesodermal cells expressing *gli3*, a transcription factor in the Hedgehog signaling pathway, contribute to both dermal and endochondral bones. We show that Gli3 regulates expression of *activin A receptor type 1-like*, a BMP type 1 receptor lost in tetrapod lineages, and thereby determines endochondral and dermal ossification. Intriguingly, Gli and Hedgehog compound knockout fish exhibited an unexpected combination of actinopterygian fish and stem-tetrapod pectoral girdle characteristics. These ontogenetic and anatomical data suggest that a trade-off between the two distinct ossification pathways is a deeply embedded developmental program in bony fishes, with potential for tuning of this trade-off to generate novel pectoral girdle forms akin to stem-tetrapods at the dawn of vertebrate terrestrialization.

---

Across the fish-to-tetrapod transition, vertebrate bones throughout the body were reconfigured from dermal (direct ossification) to endochondral bones (ossification replacing a cartilage template), facilitating terrestrial feeding, locomotion, and breathing^1,6–8^. The pectoral/shoulder girdle is a dramatic example of this dermal-to-endochondral skeletal shift. In ray-finned (actinopterygian) and lobe-finned (sarcopterygian) fishes, the pectoral girdle consists of a relatively small endochondral component (scapula and coracoid bones) and a large series of dermal bones (i.e., cleithrum, clavicle, and supracleithrum) that link the scapula and coracoid to the skull^9^ (Fig. 1a). As vertebrates transitioned onto land in the Late Devonian, the scapula and coracoid expanded, serving as robust attachment sites of forelimb musculature for terrestrial locomotion^2,3,10^. Concomitantly, a reduction of the dermal series disconnected the pectoral girdle from the skull and triggered the evolution of a functional neck, providing greater mobility to the head^11^. Despite the functional implications of the dermal-to-endochondral shift in pectoral girdle evolution, the underlying genetic and developmental changes remain unexplored.

**Figure 1.**
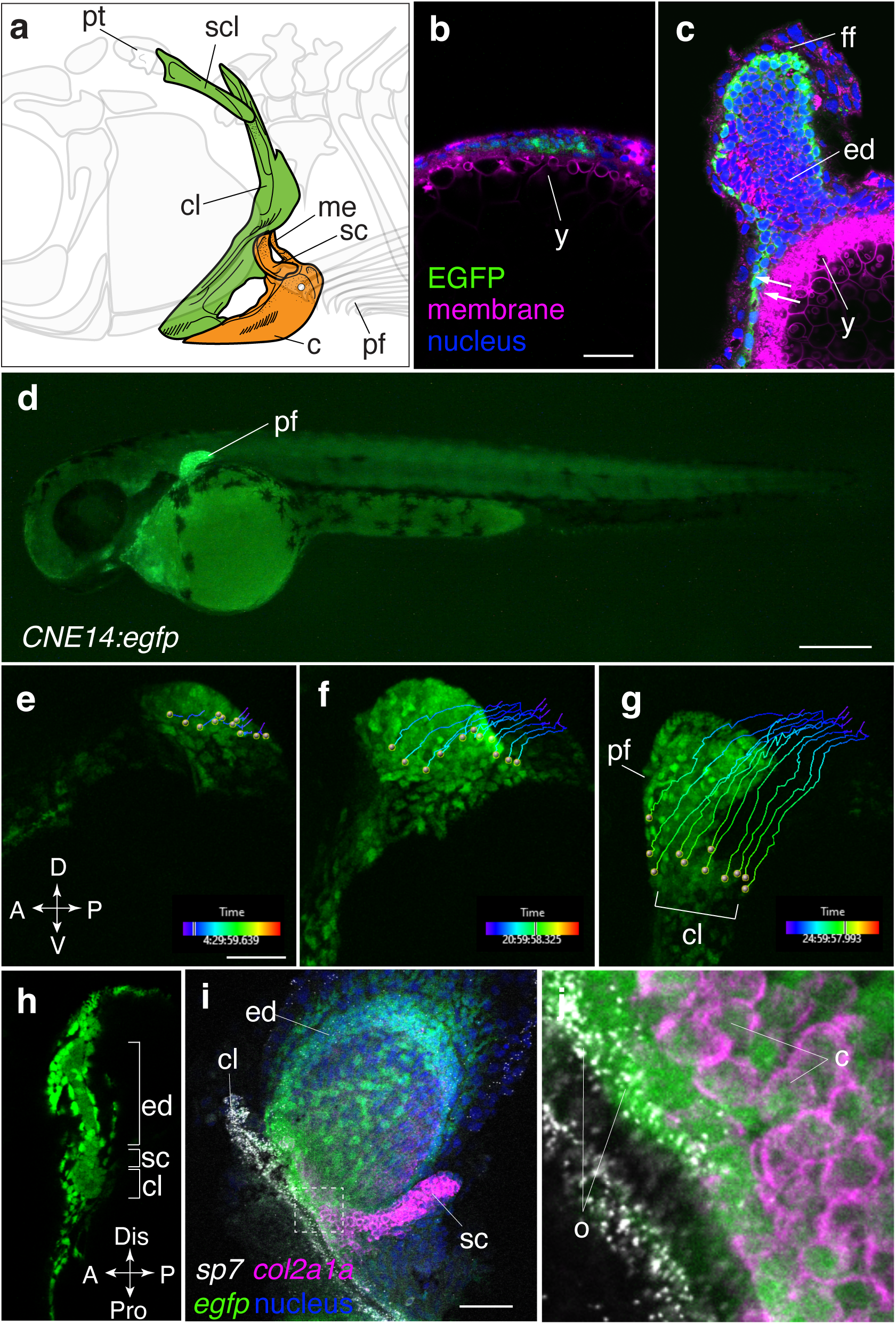
*Gli3*-positive cells contribute to the cleithrum and the scapulocoracoid in the pectoral girdle. **a**) The adult zebrafish pectoral girdle bones. The dermal bones and endochondral bones are highlighted by green and orange, respectively. The dermal series are connected to the posterior skull via the post-temporal bone (pt). The pectoral fin articulates to the scapula (sc). **b** and **c**) Digitally sectioned immunofluorescence of EGFP with membrane (CellMask) and nucleus (DAPI) staining at 32 hpf (**b**) and 48 hpf (**c**). Bright EGFP expression is observed in the mesenchymal cells at the prospective pectoral fin region at 32 hpf. As the pectoral fin grows, EGFP-positive cells are localized in the fin mesenchyme, including the proximal fin, at 48 hpf. These images were 3D-digital sections obtained by a confocal microscope. **d**) A stereotype fluorescent microscope image of *CNE14:egfp* transgenic embryo at 48 hpf. The transgenic embryos exhibit bright EGFP expression in the pectoral fin. **e-h**) Confocal microscope maximum intensity projections of live fluorescent imaging of the pectoral fin in *CNE14:egfp* embryo from 32 (**e**) to 48.5 (**f**) and to 52.5 hpf (**g**). At 32 hpf, the prospective scapulocoracoid and cleithrum cells reside inside or proximal to the pectoral fin (yellow dots in **e**). These cells migrate ventrally (**f**) and form pectoral girdle bones as development proceeds (**g**). Cell migration trajectories are dragon tails created by Imaris. The unit of the time bars is hours. *n*=16. A; anterior, P; posterior, D; dorsal, and V; ventral. **h**) an example of the original scanned image of the live fluorescent imaging data. EGFP-positive cells contribute to the endochondral disc (ed), the scapulocoracoid (sc), and the cleithrum (cl). A; anterior, P; posterior, D; dorsal, and Pro; proximal. **i** and **j**) HCR of *col2a1a* for chondrocytes (scapulocoracoid, magenta), *sp7* for osteoblasts (cleithrum, white), and *egfp* (*gli3*-positive cells, green) with DAPI staining (nuclei, blue) at 72 hpf. **j**) the enlarged image of the proximal pectoral fin marked by the rectangle in **i**. *Egfp*-positive cells express *col2a1a* in the scapulocoracoid and *sp7* in the cleithrum, indicating that *gli3*-positive cells differentiate into proliferating chondrocytes and osteoblasts. *n*= 23. Scale bars are 20 μm (**b**), 300 μm (**d**), and 50 μm (**e** and **i**). pf; pectoral fin, y; yolk, ed; endochondral disc, ff; fin fold, cl; cleithrum, sc; scapulocoracoid, o; osteoblasts, and c; chondrocytes.

The entrenched idea in the field has been that the ontogenetic sources of the two bone types are distinct cell populations; dermal bones have been proposed to develop from neural crest cells, whereas endochondral bones are thought to develop from mesodermal cells^12^. However, recent studies have provided some evidence that the two distinct ossification modes may share a common developmental origin in tetrapods^13–15^. For example, in chicken and mouse embryos the clavicle develops via dermal ossification from the lateral plate mesoderm (LPM) cells^16–19^. On the other hand, the endochondral scapula originates as an admixture of the LPM and the paraxial mesoderm (PAM) cells in mice, chicken, axolotls, and tentatively turtles^20–25^. A previous study in mice suggested that neural crest cells also contribute to the muscle attachment site of the scapula^26^. Assuming that the cleithrum in actinopterygian fishes has a neural crest origin, this study hypothesized that the neural crest contribution to the scapula is a remnant of the lost cleithrum - “the cleithrum’s ghost.” The neural crest contribution to the scapula, however, has not been discerned in other studies in tetrapods^26–28^, and thus is still under debate. Moreover, the developmental origins of the pectoral girdle bones in vertebrates with paired fins (chondrichthyan, actinopterygian, and sarcopterygian fishes except for tetrapods) remain unidentified^29^.

The insufficient knowledge about the genetic pathways responsible for piscine pectoral girdle formation is a major obstacle to characterizing the molecular machinery responsible for the evolution of the terrestrial pectoral girdle over the course of the fish-to-tetrapod transition. In mice, the *pbx* genes, which encode TALE homeoprotein transcription factors, play pivotal roles in endochondral acromion and scapular blade formation, inducing the expression of various transcription factors, such as *alx1*, *tbx15*, *gli3*, and *pax1*^30–33^. Additionally, *tbx5^−/−^* mice exhibit a loss of the entire scapula^34^. In contrast to the accumulated knowledge of the genetic mechanisms required for mammalian and avian pectoral girdle formation, the knowledge of genes involved in fish pectoral girdle formation is limited to a few studies^35–38^.

Previous studies have established that Gli3, Gli family zinc finger protein 3, plays a key regulatory role in shoulder girdle formation as well as in the patterning of distal appendages in tetrapods^39–41^, therefore, *gli3* is a compelling candidate gene to dissect the developmental mechanisms of the pectoral girdle in vertebrates with paired fins. Mechanistically, Gli3 is a component of the Hedgehog (Hh) - Gli signaling pathway, where it functions as a transcriptional activator when Hh ligands bind to a membrane receptor Patched (Ptch) or undergoes proteolytic cleavage to a transcriptional repressor without Hh ligands^42^. In tetrapod limb development, *gli3* is expressed in the anterior limb bud compared to the posteriorly localized *sonic hedgehog* (*shh*) expression, and thus is mostly processed to the repressor form without Hh ligands^40^. Consistently, *gli3* deletion in mice causes a wide scapula blade and polydactyly, demonstrating that the Gli3 repressor restricts scapula width and digit number^31,40,43,44^. Deletion of *gli3* in the actinopterygian medaka has further supported the notion that it has an evolutionary conserved function in repressing fin ray and endochondral bone number in the pectoral fin^45^. Moreover, the previous studies demonstrated that *shha* (one of the shh genes created in the third-round whole genome duplication) is indispensable for the formation of the cleithrum and the scapulocoracoid, a primordium of the scapula and coracoid, in the zebrafish and medaka pectoral girdle^46,47^. However, further investigation of Hh-Gli signaling is crucial to elucidate the genetic mechanisms of dermal and endochondral bone specification and development in vertebrates with paired fins.

To identify the embryonic origins of the two types of ossification in the pectoral girdle of aquatic fish, we genetically mapped the contribution of *gli3*-positive cells to pectoral girdle bones using the zebrafish transgenic system. *Gli3*-positive cells are of lateral plate mesoderm origin in tetrapods^44,48–50^. We cloned the conserved noncoding element 14 (*CNE14*), an evolutionarily conserved *gli3* enhancer for pectoral appendage expression^44^, from the chondrichthyan elephant sharks. Chondrichthyans have not undergone the teleost fish-specific third round whole genome duplication and thus likely retain evolutionarily conserved activities of cis-regulatory elements^51^. Next, we established *CNE14:egfp* transgenic zebrafish, which show EGFP expression in the fin mesenchyme from 32 hours post fertilization (hpf) and beyond (Fig. 1b-d, Extended Data Table 1). Fluorescent live-cell imaging of *CNE14:egfp* in the pectoral fin from 32 to 72 hpf showed that EGFP-positive cells inside or peripheral to the proximal pectoral fin at 32 hpf migrate in the distal to proximal direction and contribute to the cleithrum and scapulocoracoid at 72 hpf (Fig. 1e-h). Multiplexed fluorescent *in situ* hybridization chain reaction (HCR) with probes complementary to the osteoblast marker *sp7*^52^, the proliferative chondrocyte marker *col2a1a*^53^, and *egfp* (*gli3*-positive cells) demonstrated that these EGFP-positive cells differentiate to osteoblasts in the cleithrum and to chondrocytes in the scapulocoracoid (Fig. 1i and j). Taken together, these results demonstrate that *gli3*-positive cells inside and peripheral to the early fin bud migrate and differentiate to osteoblasts in the cleithrum or chondrocytes in the scapulocoracoid in zebrafish embryos.

To test the function of Gli3 in dermal and endochondral ossification in zebrafish, we generated stable *gli3* homozygous knockout zebrafish mutants bearing a frame-shift mutation in exon5 (*gli3*^14ins/14ins^) using CRISPR/Cas9 (Extended data Fig. 1, 2, and Extended Data Table 1). HCR with probes complementary to *sp7* and *col2a1a* showed that fewer osteoblasts are present in the cleithrum at the level of the scapulocoracoid in *gli3*^14ins/14ins^ mutants compared to wildtype embryos at 72 hpf (Fig. 2a and b). Gli3 functions as a transcriptional activator or repressor depending on the presence of Hh^28^. Therefore, osteoblast reduction could arise from a loss of either or both functions. To differentiate between these possibilities, we treated wildtype zebrafish embryos with BMS-833923, a Smoothened antagonist / Hedgehog inhibitor^54^, which decreases the amount of Gli activator, from 32-72 hpf. In BMS-833923 treated embryos, we observed fewer osteoblasts in the cleithrum at the level of the scapulocoracoid in a comparable manner to *gli3*^14ins/14ins^ embryos at 72 hpf (Extended Data Fig. 3). This demonstrates the functional significance of Gli as an activator in osteoblast differentiation.

**Figure 2.**
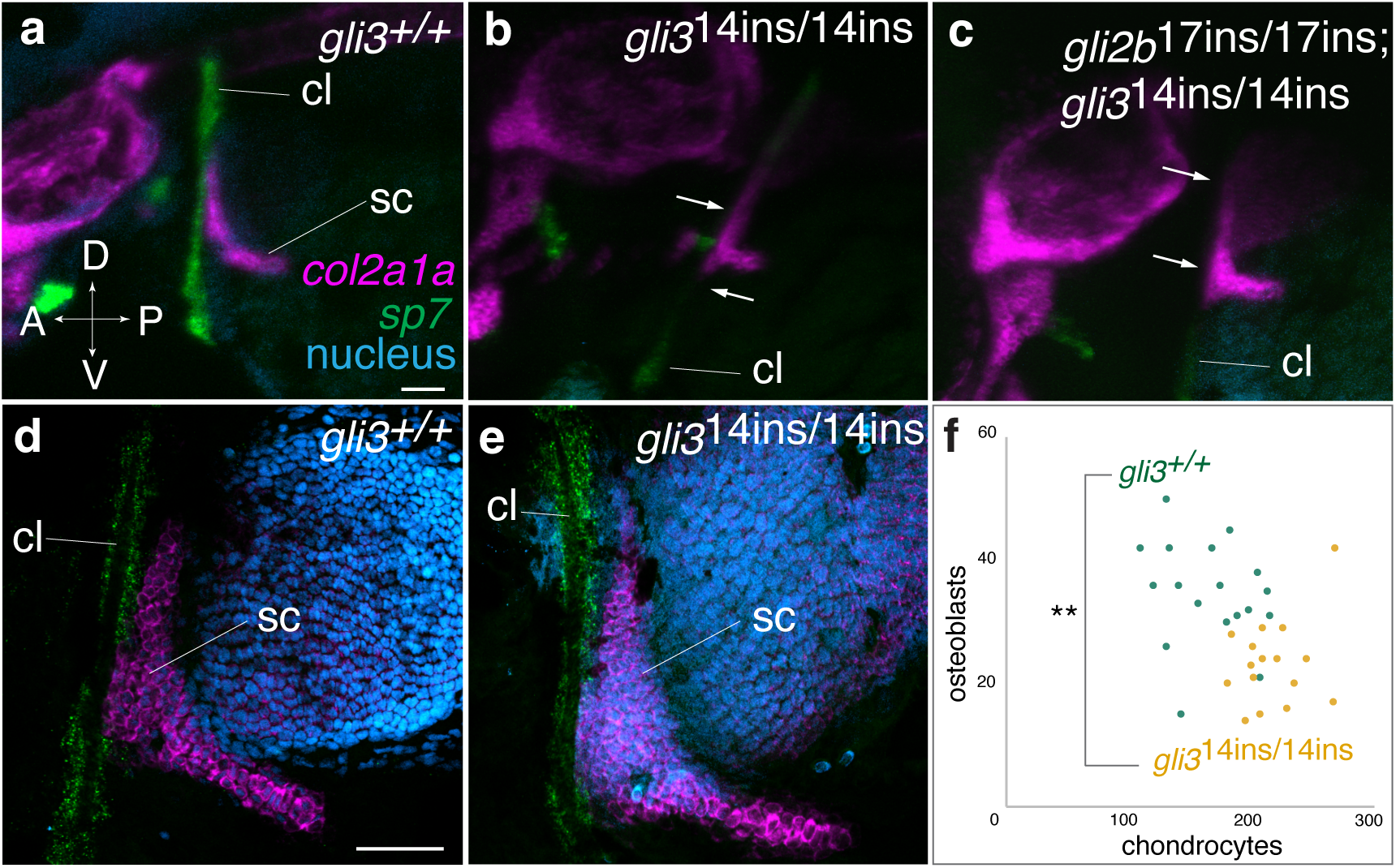
The trade-off of osteoblast and chondrocyte differentiation regulated by *gli* genes. **a-c**) HCR of *col2a1a* (magenta) and *sp7* (green) with DAPI staining (cyan). 3D stack lateral visualization of confocal scanning of *gli3*^+/+^ (**a**), *gli3*^14ins/14ins^ (**b**), and *gli2*^17del/17del^ *; gli3*^14ins/14ins^ embryos (**c**) or of dissected pectoral fins of *gli3*^+/+^ (**d**) and *gli3*^14ins/14ins^ (**e**) embryos at 72 hpf. The osteoblasts around the cleithrum at the level of the scapulocoracoid are reduced in *gli3*^14ins/14ins^ and *gli2*^17del/17del^ *; gli3*^14ins/14ins^ embryos (**b** and **c**). *n*= 6 for each genotype in **a**-**c**. The scapulocoracoid is larger in *gli3*^14ins/14ins^ embryos (**e**) than in *gli3*^+/+^ embryos (**d**). **f**). The comparison of the osteoblast (*sp7*-positive) and chondrocyte (*col2a1aa*-positive) numbers in the cleithrum and the scapulocoracoid, respectively, between *gli3*^+/+^ and *gli3*^14ins/14ins^ embryos (*n*=18 for *gli3*^+/+^ and *n*=16 for *gli3*^14ins/14ins^ embryos). **; *p*<0.01 by a student t-test. Scale bars are 300 μm in **a** and 50 μm in **d**. cl; cleithrum, sc; scapulocoracoid.

**Figure 3.**
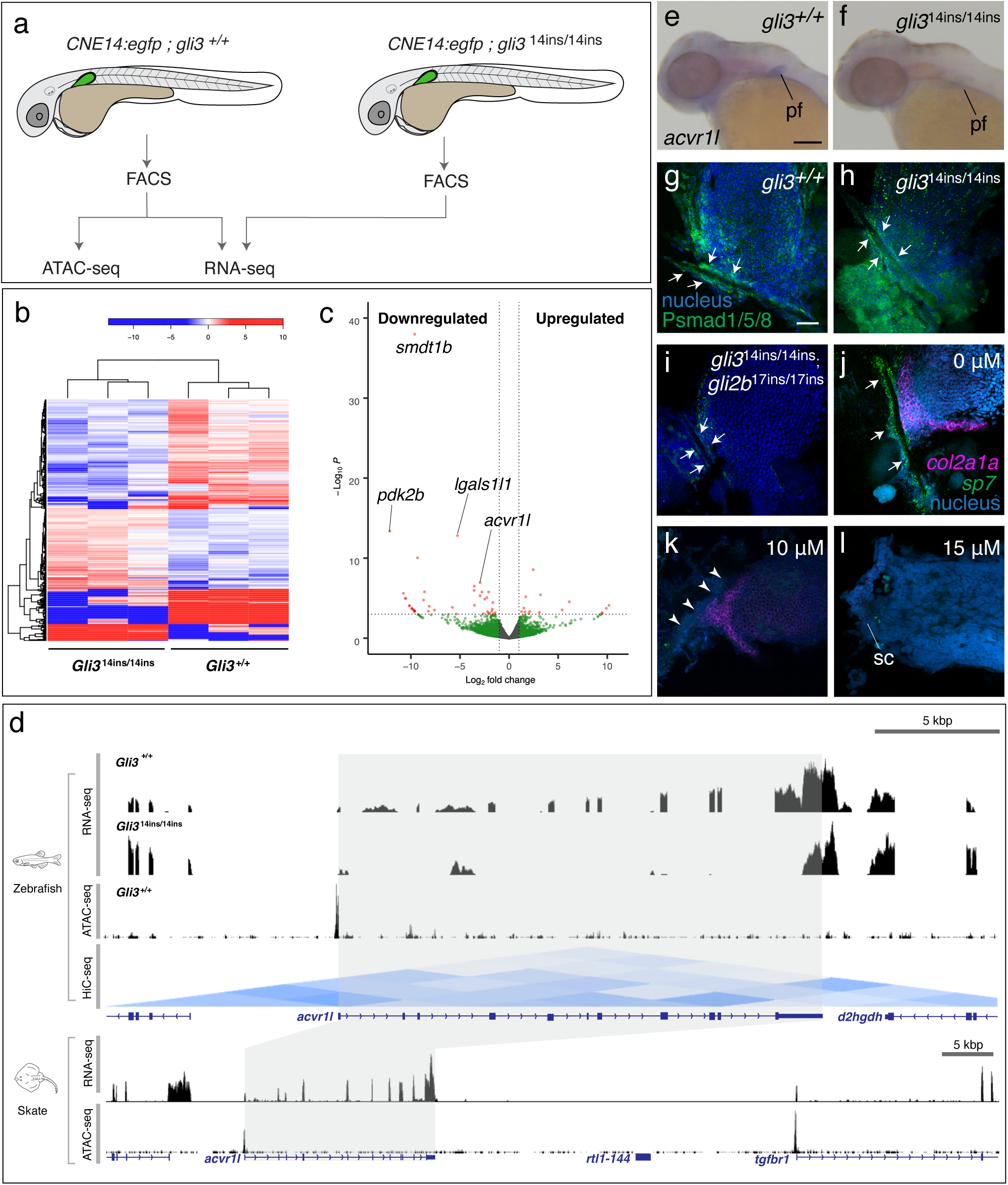
Gli genes regulate BMP signaling via *activin receptor type 1-like* expression. **a)**The experimental scheme of the unbiased high-throughput genomic screening (see Methods). From *CNE14:egfp; gli3*^+/+^ and *CNE14:egfp; gli3*^14ins/14ins^ embryos, EGFP-positive cells were separately isolated by FACS. The sorted cells were subjected to RNA-seq and ATAC-seq. Three biological replicates were conducted for RNA-sequencing and two were for ATAC-seq. **b)** The heat map of top 2,000 differentially expressed genes in *CNE14:egfp; gli3*^+/+^ and *CNE14:egfp; gli3*^14ins/14ins^ embryos. Color code; red indicates high and blue indicates low enrichment. **c)** The volcano plot of the RNA-seq result. The genes with p<0.5 and >two-fold expression change are coded by red. *Acvr11* is one of the prominent Gli3 candidate genes. **d**) the genome browser visualization of zebrafish RNA-seq (wildtype and *gli3*^14ins/14ins^ samples), ATAC-seq, and HiC results and skate RNA-seq and ATAC-seq results at the *acvr1l* locus. In zebrafish, the transcript level is lower at the exons of *acvr1l* in *gli3*^14ins/14ins^ embryos than *gli3*^+/+^ embryos. Zebrafish ATAC-seq and HiC showed that the promoter region of *acvr1l* is ACR and in a TAD. Zebrafish HiC result was produced from the previously published data^62^. Skate RNA-seq and ATAC-seq was generated from the previous paper^77^. **e, f**) The expression pattern of *acvr1l* in *gli3*^+/+^ and *gli3*^14ins/14ins^ embryos. Expression in the pectoral fin is decreased in *gli3*^14ins/14ins^ embryos compared to *gli3*^+/+^ embryos. *n*= 24. **g** -**i**) Immunofluorescence of phosphorylated-Smad 1/5/8. The strong signal is observed in cells surrounding the cleithrum of wildtype embryos (**g, arrows**), which decreases to *gli3*^14ins/14ins^ (**h**) and to *gli2b*^17ins/17ins^*; gli3*^14ins/14ins^ embryos (**i**). *n*= 7 for each genotype. **j-l**) LDN193189-treated pectoral fins stained by HCR with *sp7* and *col2a1a* probes and DAPI staining. *Sp7* expression was diminished by 10 μM of the inhibitor treatment and both *sp7* and *col2a1a* were not detected in 15 μM treatment. 15 μM LDN193189-treated embryos also showed a loss of the cleithrum. Arrows indicate *sp7* expression and arrowheads show its weak or no expression. sc; scapula. *n*= 23 for 0 and 10 μM and 8 for 15 μM treatment. The scale bar in **e** and **g** are 50 μm. **g**-**l** are the same scale.

To genetically corroborate that Gli functions as an activator in the zebrafish pectoral girdle, we decreased the amount of Gli activator by introducing a stable frameshift mutation into the coding sequence of *gli2b*, another *gli* family zinc finger protein that functions as a transcriptional activator^55^, in *gli3*^14ins/14ins^ embryos using CRISPR/Cas9 (Extended Data Fig. 1, Extended Data Table 1). Consistent with the phenotype of BMS-833923-treated embryos, *gli2b*^17del/17del^*; gli3*^14ins/14ins^ double knockout embryos have fewer osteoblasts in the cleithrum at the position where the scapulocoracoid attaches than *gli3*^14ins/14ins^ embryos at 72 hpf (Fig. 2c). We also observed that the number of *col2a1a*-positive proliferative chondrocytes in the scapulocoracoid of *gli3* ^14ins/14ins^ embryos increased compared to wildtype embryos (Fig.2 g-i and Extended Data Fig.3). These results demonstrate that the Gli3 activator induces osteoblast differentiation in the cleithrum at the expense of chondrocytes.

To elucidate the Gli3 pathway that determines osteoblast and chondrocyte differentiation, we identified Gil3 target genes in an unbiased way by combining RNA-sequencing (RNA-seq) and Assay for Transposase-Accessible Chromatin using sequencing (ATAC-seq) in wildtype and *gli3*^14ins/14ins^ embryos (Fig. 3a). We separately isolated EGFP-positive cells from *CNE14:egfp; gli3*^+/+^ and *CNE14:egfp; gli3*^14ins/14ins^ embryos by Fluorescence Activated Cell Sorting (FACS) and conducted RNA-seq at 55hpf, when both the cleithrum and the scapulocoracoid are at their early developmental stages (Fig 3a). A comparative analysis of transcriptome profiles between EGFP-positive cells in *gli3*^+/+^ and *gli3*^14ins/14ins^ embryos identified 351 significantly upregulated genes and 232 downregulated genes (*p*<0.05) (Fig. 3b, Extended Data Table 2). This gene list includes *klf3* and *zic3*, which were previously identified as Gli3 direct target genes in mouse limb bud^56^, supporting the efficacy of our method. Next, we conducted ATAC-seq with EGFP-positive cells sorted from *CNE14:egfp; gli3*^+/+^ embryos at 55 hpf, identified accessible chromatin regions (ACRs) in a genome-wide manner, and annotated these ACRs to genes depending on the proximity to transcription start sites (Extended Data Table 3). The gene with one of the lowest *p*-values from RNA-seq and an ACR promoter is *activin A receptor type 1-like* (*acvr1l*), which encodes a BMP receptor type 1^57^ (Fig. 3c). Given the necessity of BMP signaling in osteoblast and chondrocyte differentiation and its involvement in human skeletal diseases^58–61^, *acvr1l* is a compelling Gli3 target for pectoral girdle development. An analysis of previously published Hi-C data indicates that the *acvr1l* promoter domain is in a ∼90 kbp topologically associating domain (TAD), suggesting its cis-regulatory elements lie in this region^62^ (Fig. 3d). Intriguingly, *acvr1l* gene is evolutionarily conserved in actinopterygians, yet lost in the tetrapod lineage and has thus been implied to be a “gene left in the water” associated with the fish-to-tetrapod transition^63^.

Subsequent whole-mount RNA *in situ* hybridization corroborated the regulation of *acvr1l* by Gli3; *acvr1l* transcripts are enriched in the endochondral disc of the wildtype pectoral fin but are decreased in *gli3*^14ins/14ins^ fin at 55 hpf (Fig. 3e, f). Moreover, we found *acvr1l* expression in the pectoral fin is a shared feature with chondrichthyans (*Leucoraja erinacea*) (Extended Data Fig. 4). In accordance with the reduction of *acvr1l* expression in *gli3*^14ins/14ins^ embryos, we also found a decrease in staining levels of phosphorylated Smad1/5/8 (active forms of Smad1/5/8 in BMP signaling) in osteoblasts surrounding the cleithrum bone from wildtype to *gli3*^14ins/14ins^ and to *gli3*^14ins/14ins^; *gli2b*^17del/17del^ (Fig. 3g-i). Finally, to test the function of Acvr1l in the differentiation of osteoblasts and chondrocytes in shoulder girdle formation, we treated wildtype zebrafish embryos with an Activin receptor type 1 inhibitor LDN193189^64^, from 32 to 72 hpf at different concentrations. Compared to the proper expression of *sp7* and *col2a1a* in the control pectoral girdle, *sp7* or both *sp7* and *col2a1a* expression were diminished in 10 μM and 15 μM LDN193189-treated embryos, respectively (Fig. 3j-l). Therefore, the differentiation of osteoblasts and chondrocytes requires distinct thresholds of BMP signaling via Acvr11 in the pectoral girdle.

Survival of *gli3*^14ins/14ins^ fish to adulthood allowed us to determine the functional roles of the Hh-Gli signaling pathway in the composition of the adult pectoral girdle bones. We tested for alterations of skeletal morphology found in *gli3*^14ins/14ins^ fish using skeletal staining and µCT scanning (Methods, Extended Data Fig. 5). As compared to wildtype zebrafish, *Gli3*^14ins/14ins^ fish at three months old have excessively mineralized skull bones and ectopic ossification in the eye lens (Extended data Fig. 5, n=5/7). These phenotypes are consistent with ones observed in the human *gli3* congenital disease Greig Cephalopolysyndactyly Syndrome^56^.

The pectoral girdle in non-teleost actinopterygians consists of a series of dermal (supracleithrum, cleithrum, and clavicle) and endochondral bones (scapulocoracoid and mesocoracoid)^65^. Teleosts, including zebrafish and medaka, share these fundamental components, except for the separation of the scapula and coracoid and the loss of the clavicles^66^ (Fig. 4a). Gross examination and skeletal staining showed that in approximately 20 % of *gli3*^14ins/14ins^ fish either the left or right pectoral fin shifted dorsally due to the dorsoventrally shortened cleithrum (28/159 fish, Extended Data Figure 5, Extended Data Table 4). Intriguingly, subsequent µCT scanning and three-dimensional morphometric analysis revealed mixed characteristics of actinopterygian and stem-tetrapodomorph pectoral girdle morphologies in these severely affected girdles; the dorsal blade of the cleithrum and the posterior branchial lamina, which posteriorly supports opercular chamber movement in fish, were markedly reduced (Fig. 4a, d, k, Extended Data Figure 6). Moreover, the supracleithrum was absent (Fig. 4a, d), resembling the condition of stem-tetrapods such as *Tulerpeton*^3,4^. The cleithrum on the opposite side in these severely affected fish and on both sides in the remaining 80% of *glis3*^14ins/14ins^ zebrafish was dorsoventrally shorter compared to those of wildtype individuals (Extended data Figure 7, n=23/23). In contrast to the reduction of the dermal series, the scapula of *gli3*^14ins/14ins^ fish expanded with characteristics of stem-tetrapod traits; the enlarged supraglenoid buttress, and the anteromedially extended flange for the glenoid (Fig. 4b, e). The glenoid was ovoid and concave in *gli3*^14ins/14ins^ fish, and it is posterolaterally opened compared to the flat glenoid in wildtype fish. This is a prominent feature of stem-tetrapods, such as *Tiktaalik* or *Acanthostega,* that permits the rotation, flexion, extension, protraction, and retraction of the humerus^2^ (Fig. 4b, c, e, f; n=3/6).

**Figure 4.**
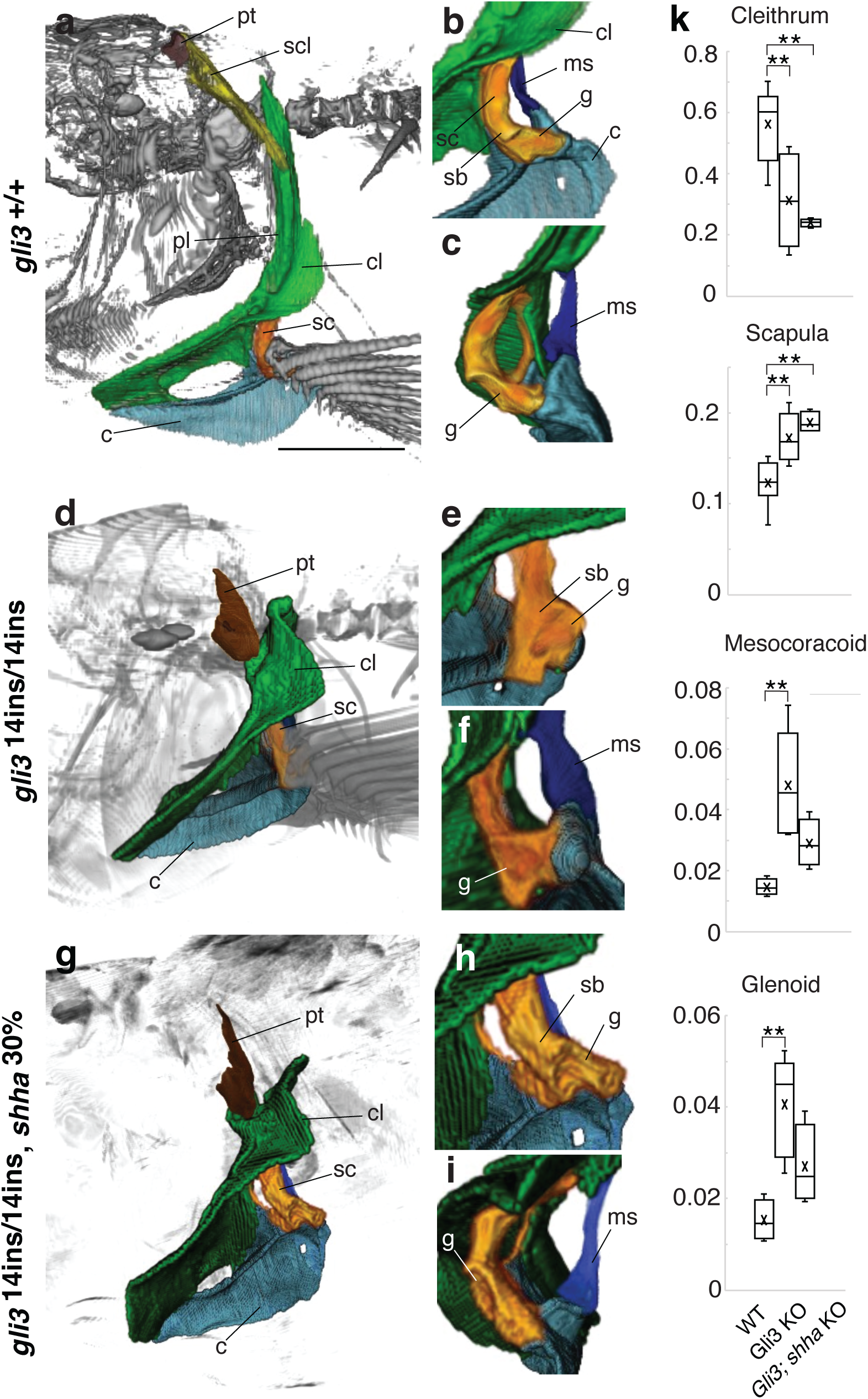
Adult pectoral girdle phenotype in Hh-gli signaling knockout fish. **(a-c, d-f, g-i) µ**CT scan and following segmentation analysis of three-months old adult fish. **a-c)** wildtype, **d-f)** *gli3*^14ins/14ins^, **g-i**) *gli3*^14ins/14ins^*; shha*^30%KO^ fish. **a, d, g**) the whole-body 3D reconstruction, **b, c, e, f, h, i**) the enlarged images of the scapulocoracoid and mesocoracoid (**b**, **e**, **h**; lateral view, **c**, **f**, **i**; posterior view). In contrast to the thin and long dorsal cleithrum with the postbranchial lamina in *gli3*^+/+^ fish (**a**), the dorsal cleithrum was reduced in either the left of right side of *gli3*^14ins/14ins^ fish (n=28/159) (**d**) and both sides of *gli3*^14ins/14ins^*; shha*^30%KO^ fish (3/3) (**g**). In the scapula of *gli3*^+/+^ fish, the glenoid is anteroposteriorly wide and flat (**b, c**), but it is concave and posterolaterally opened, and delineates the proximal surface of basal radials of the pectoral fin in *gli3*^14ins/14ins^ and *gli3*^14ins/14ins^*; shha*^30%KO^ fish (**e, f, h, i**) (n=5/6 for each genotype). The supraglenoid buttress is more developed in *gli3*^14ins/14ins^ and *gli3*^14ins/14ins^*; shha*^30%KO^ fish compared to wildtype fish (**b**, **e**, **h**). **k**) Quantitative analysis of the cleithrum, scapula, coracoid, and glenoid size. The length and width of each bone were measured and standardized by the dorsoventral length of the skull (see Methods). While the cleithrum is dorsoventrally shorter in *gli3*^14ins/14ins^ and *gli3*^14ins/14ins^*; shha*^30%KO^ fish than *gli3*^+/+^ fish, the scapula and mesocoracoid is larger in *gli3*^14ins/14ins^ and *gli3*^14ins/14ins^*; shha*^30%KO^ fish. The glenoid is more concave in *gli3*^14ins/14ins^ and *gli3*^14ins/14ins^*; shha*^30%KO^ fish than wildtype fish. ANNOVA test and following Tukey-Kramer Post Hoc test were conducted to detect statistically significant differences of bone size among wildtype and knockout fish. **; absolute mean value > Q critical value in Tukey-Kramer Post Hoc Test. 5.07 and 5.83 > 3.73 in the cleithrum, 5.10 and 6.16 > 3.73 in the scapula, and 6.07 > 3.88 in the mesocoracoid, and 6.05 > 3.88 in the glenoid). Maximum, minimum, median, and average (“X”) values are shown in the box and whisker plots. c; coracoid, cl; cleithrum, g; glenoid, ms; mesocoracoid, pl; postbranchial lamina, sb; supraglenoid buttress, and scl; supracleithrum. The scale bar indicates 2.5 mm.

**Figure 5.**
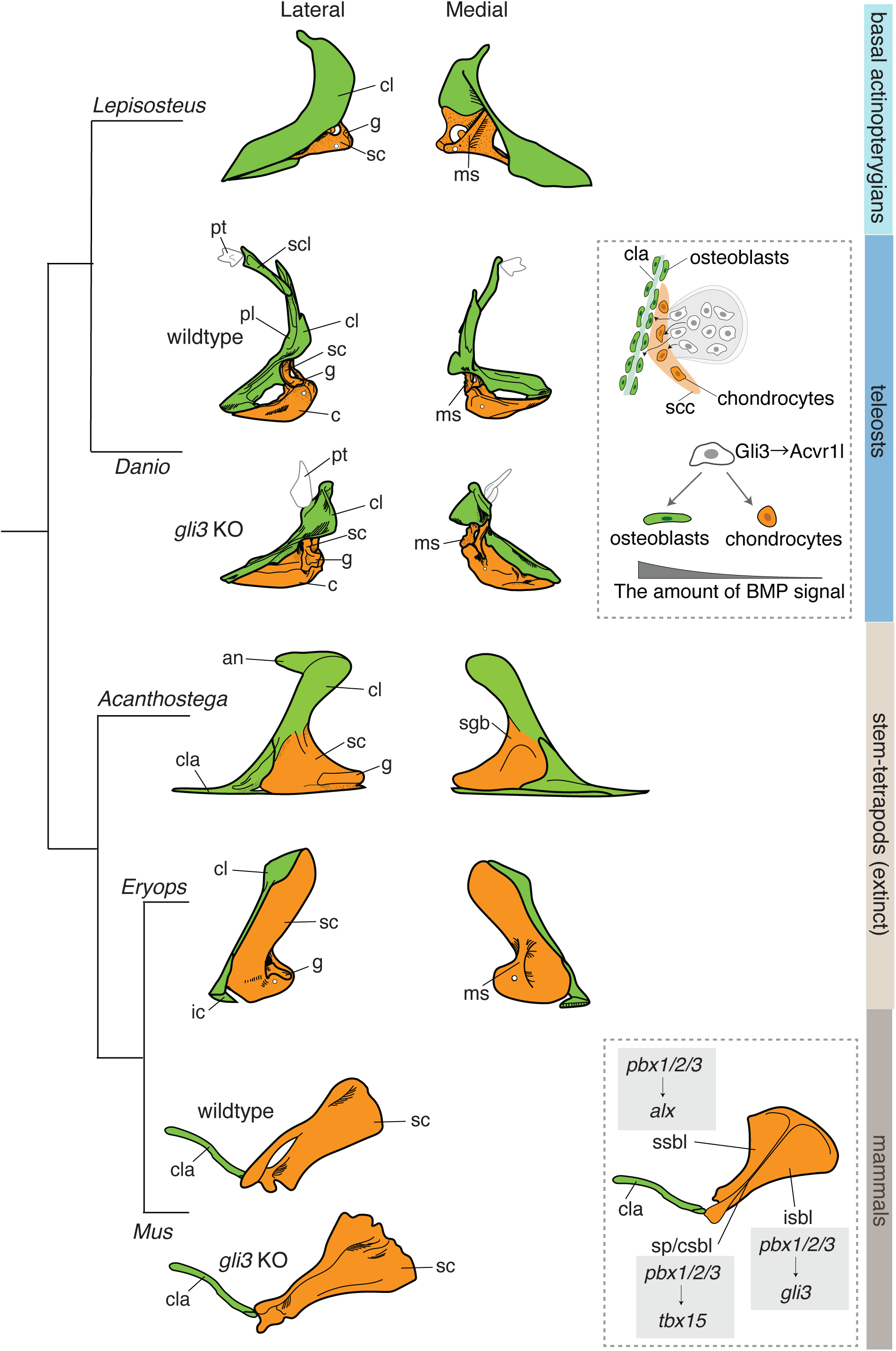
Pectoral girdle evolution from fish to tetrapods and Gli3 function in their development. From fish to tetrapods, endochondral bones (scapula and coracoid) expanded at the expense of the dermal bones (anocleithrum, cleithrum, and clavicles). In *gli3*^14ins/14ins^ zebrafish, the cleithrum becomes dorsoventrally short, reminiscent of the pectoral girdles of early tetrapods, such as *Acanthostega*, which were evolving to live on land. *Gli3*-positive cells migrate into the cleithrum and scapulocoracoid, and Gli3 regulates BMP signaling via *acvr11* expression. In crown-group tetrapods, including mice and chickens, the prospective pectoral girdle cells reside in the proximal region of forelimb buds. Pbx1/2/3 family genes regulate the expression of *alx, tbx15,* and *gli3* in the superior scapula blade (ssbl), inferior scapula blade (isbl), and spine/central scapula blade (sp/csbl) ^31,78^. Abbreviations: an, anocleithrum; c, coracoid; cla, clavicle; cl, cleithrum; g, glenoid; ic, interclavicle; ms, mesocoracoid arch; pl, postbranchial lamina; sc, scapula; scc, scapulocoracoid; scl, supracleithrum; sgb, supraglenoid buttress. The pectoral/shoulder girdles of *Lepisosteus*, *Acanthostega*, *Eryops*, and *Mus* were reproduced from previous studies with permission ^3,31,68^. The photos of the pectoral girdle of an adult *Lepisosteus oculatus* are in Extended Data Fig. 9.

The mesocoracoid is another endochondral bone that bridges the coracoid and the cleithrum and present in actinopterygian fishes but not in sarcopterygian fishes and stem-tetrapods such as *Tiktaalik* or *Acanthostega*. In these stem-tetrapods, the supraglenoid buttress is the evolutionary counterpart of the mesocoracoid^67^. The mesocoracoid/supraglenoid buttress becomes stouter from fishes with paired fins to stem-tetrapods to serve as an attachment site for the dorsomedial musculature of the forelimb^68^. In *gli3*^14ins/14ins^ zebrafish, the mesocoracoid is laterally wider than in the wildtype, which is consistent with the enlargement of the endochondral bones in the absence of Gli3 function (Fig. 4c, f, n=4/5).

Additionally, to investigate whether the pectoral girdle phenotype is exacerbated by a further reduction of Gli activator, we injected gRNA complementary to zebrafish *shha* with *Cas9* mRNA into *gli3*^14ins/14ins^ eggs, raised them for three-months, and genotyped them by deep sequencing of *shha* (see Methods). Strikingly, *gli3*^14ins/14ins^*; shha*^30%^ fish (30% of the intact *shha* is detected) exhibited a dorsoventrally short cleithrum comparable to the severely affected cleithrum in *gli3*^14ins/14ins^ fish, but on both the left and right sides (Fig.4g, n=3/3). The shape of the scapula in this compound knockout fish is also concave, posterolaterally opened, and even larger than that of wildtype and *gli3*^14ins/14ins^ fish (Fig.4h, i).

The evolutionary trajectories of the intricate shoulder girdle development during the origin of stem-tetrapods have remained uncharacterized. Taking advantage of the ontogenetic and genetic accessibility of zebrafish, we discovered that *gli3*-positive mesodermal cells give rise to both dermal and endochondral bones in the pectoral girdle. Our results show that modulations in Hh-Gli signaling are plausible evolutionary changes to assemble the tetrapodmorph pectoral girdle through mesodermal cell fate alternations, which have been also reported in the evolution of digits from fin rays^13,14^. An evolutionary novelty through a cell fate change of mesodermal cells is thus pervasive in appendage evolution across the fish-to-tetrapod transition.

The phenotype of *gli3* knockout zebrafish differs from the previously published medaka phenotype^45^, possibly reflecting diverse function of Hh-Gli signaling in the formation of exceptionally diverse pectoral girdle in actinopterygian fishes, such as the short cleithrum in medaka^69^. The dorsoventrally short cleithrum, the loss of the supracleithrum, the posterolaterally oriented concave glenoid, and the robust mesocoracoid in *gli3* single and *gli3;shh* compound knockout zebrafish conform to the shared features of the pectoral girdle bones in Devonian-era stem-tetrapods, such as *Tiktaalik*, *Acanthostega,* and *Ichthoystega*^2,3,10^. At the origin of terrestrial vertebrates, genetic alterations in Hh-Gli or associated signaling pathways, such as the evolutionary reduction and eventual loss of *acvr1l*, might have released a deeply embedded developmental program to generate tetrapodmorph pectoral girdle phenotypes in bony vertebrates.

## Methods

### Animal maintenance

All fish work was conducted according to standard protocols approved by the animal committees of Rutgers University (protocol number 201702646) and Michigan State University (PROTO202200367).

### Establishment of *CNE14:egfp* transgenic fish

The region of the elephant shark genome homologous to human CNE14^26^ was amplified by PCR and subcloned pCR™8/GW/TOPO® (Invitrogen) (the primer sequences are in Extended Data Table 1). The cloned CNE14 fragment was transferred into pXIG-cfos-EGFP vector by Gateway™ LR Clonase™ II (Invitrogen)^46^. CNE14 / pXIG-cofs-EGFP vector was injected into one-cell stage zebrafish eggs with Tol2 mRNA as previously described^47^. F0 embryos with EGFP expression were raised for three months and the EGFP expression pattern was confirmed in F1 embryos obtained from outcrossing the F0 to wildtype fish under a stereotype fluorescent microscope. The embryos with EGFP expression in the pectoral fin were reared to adult stage and further crossed to wildtype fish to establish stable transgenic embryos, which were used for immunofluorescence, HCR in situ hybridization, and live fluorescent imaging.

### Whole mount *in situ* hybridization

The 3’UTR or coding sequence of the genes analyzed in this paper were amplified from 48 hpf zebrafish cDNA by PCR and cloned into pCR™II-TOPO® vector (Invitrogen) (PCR primer sequences are in Extended Data Table 1). The RNA probes were synthesized from the vectors by T7 or SP6 RNA polymerase. Chromogenic *in situ* hybridization for *acvr1l* in zebrafish, skate, and gar embryos was performed as previously described^48^. In situ hybridization chain reaction (HCR) was performed as previously described^49^ using complementary probes to zebrafish *sp7*, *col2a1aa*, and *egfp,* which were all synthesized by Molecular Instruments (CA). To observe the HCR results at the single cell resolution, the pectoral fin and girdle complex was manually dissected out from the body with tweezers, mounted on slide glasses, and scanned by a confocal microscope (Zeiss LSM510 META). The numbers of chondrocytes in the scapulocoracoid (*col2a1a*+) and osteoblasts surrounding the cleithrum at the level of the scapulocoracoid (*sp7*+) were manually counted and statistically compared among different genotypes or embryos treated by different concentration of the inhibitor by a Student’s t-test.

### Live fluorescence cell imaging

*CNE14:egfp* transgenic fish were crossed to wildtype fish and fertilized eggs were collected. Embryos with EGFP expression in the pectoral fin primordium were selected under a fluorescent stereotype microscope at 32 hpf and embedded in 1% low-melt agarose gel in a glass-bottom 3.5 cm dish^50^. The gel covering the anterior half of embryos was removed to allow normal growth of the embryos. E3 medium including tricaine at 0.003% was added to the dishes and the dishes were kept inside an incubator chamber at 28°C by a confocal microscope (Zeiss LSM510) from 32-72 hpf. Z-stack images (3um z-interval, approximately 30-40 slices for each embryo) were obtained at 30 minutes time intervals. The obtained time lapse data was analyzed by Imaris to manually track EGFP-positive cell movement in the pectoral fin and girdle.

### Establishment of *gli3*, *gli2b;gli3*, and *gli3;shh* compound knockout fish

*Gli3* single knockout fish with frameshift mutations in exon5 and exon14 were generated by injection of gRNAs complementary to target sites and *Cas9* mRNA into wildtype (*AB) fertilized eggs as previously reported (Nakamura 2016). The detailed information of frame shift mutations is summarized in Extended Data Figure 1. Briefly, gRNA and *Cas9* mRNA were injected into zebrafish one-cell stage eggs, and the injected eggs were reared to adult fish, which were subjected to T7 assay and sequencing to determine genetic mutations at the target loci. The fish with frame-shift mutations (F0) were out-crossed to wildtype fish to obtain heterozygous knockout fish (F1). To avoid any off-target effects of CRISPR/Cas9 gene deletions on the phenotypic analysis, we repeated outcrossing the heterozygous knockout fish to wildtype fish (*AB) and raising offspring twice, obtaining F4 fish for the analysis conducted in the paper. Then, *gli3 ex5*^14ins/+^ fish (F4) were crossed with each other and *gli3 ex5*^14ins/14ins^ fish were established. The survival rate of *gli3 ex5*^14ins/14ins^ zebrafish is not significantly different from that of wildtype fish and the ratio of *gli3* ^+/+,^ *gli3 ex5*^14ins/+^, and *gli3 ex5*^14ins/14ins^ follow Mendelian inheritance ratio. To obtain *gli2/gli3* compound knockout fish, gRNA complementary to a target site in *gli2b* exon1 and *Cas9* mRNA was injected into one-cell stage *gli3 ex5*^14ins/14ins^ fertilized eggs. The injected eggs were reared to an adult stage and subjected to a T7 assay and sequencing to identify frameshift mutations. The fish with frameshift mutations were outcrossed to *gli3 ex5*^14ins/14ins^ fish and the collected eggs were raised for three months and genotyped for *gli2b*. Obtained *gli2b*^17del/+^*; gli3 ex5*^14ins/14ins^ fish were crossed with each other and embryos were subjected to HCR *in situ* hybridization.

To obtain *gli3/shha* compound knockout fish, gRNA complimentary to *shha* and *Cas9* mRNA were injected into *gli3 ex5*^14ins/14ins^ fertilized eggs. The tail fins of three-month old adult fish were cut and subjected to a T7 assay for the *shha* gene. The tail lysis of the mutant fish was further subjected to deep sequencing of *shha* target loci to determine frameshift mutation ratio among all mutations (Extended Data Table 1).

All gRNA sequence and genotyping primers are summarized in Extended Data Table1. The frameshift mutations in all mutant fish are in Extended Data Figure 1.

### Genotyping of *gli3* and *gli2b* knockout fish

Adult or embryonic tails were excised and lysed as previously described^14^. Using these tail lysis solutions as PCR templates, the *gli3* and *gli2* loci were amplified by PCR respectively. The PCR primers used for PCR amplification are summarized in Extended Data Table 1. For *gli3*, a 14 bp difference between wildtype and mutant fish was confirmed on 3% agarose gels. BSAJ1 was added to *gli2b* PCR products and the treated PCR products were confirmed on 1% agarose gel (wildtype; 232 bp, *gli2b*^17ins^; 130 + 123 bp).

### Chemical inhibitor treatment of zebrafish embryos

Zebrafish embryos were cultured in 40 ml of E3 medium (5 mM NaCl, 0.17 mM KCl, 0.33 mM CaCl_2_, 0.33 mM MgSO_4_) containing Smoothened inhibitor (BMS-833923, Selleckchem) or Activin receptor type 1 inhibitor (LDN 193189, TOCRIS) in 10 cm plastic dishes from 32 hpf to 72 hpf. The concentration of BMS-833923 was 5 μM and of LDN 193189 was 0, 10, and 15 μM. At 72 hpf, these inhibitor-treated embryos were fixed by 4% PFA and subjected to immunofluorescence staining or HCR. The 15 μM LDN 193189-treated embryos showed high mortality (12/20 embryos), and the only surviving embryos were used for HCR ISH and subsequent confocal scanning.

### FACS sorting of EGFP-positive cells from *CNE14;egfp* fish

Approximately 100,000 EGFP-positive cells were collected from *CNE14:egfp; gli3*^+/+^ and *gli3*^14ins/14ins^ fish by FACS as previously described^51^. Briefly, *CNE14:egfp; gli3*^+/+^ or *gli3*^14ins/14ins^ fish were crossed to wildtype fish or and *gli3*^14ins/14ins^ fish, respectively, and fertilized eggs were collected. The eggs were raised to 55 hpf at 27.5°C. Approximately 200 embryos from each genotype were separately dissociated into single cells. The yolk was removed by de-yolk buffer and embryos were rinsed by PBS. Then, embryos were dissociated into single cells by 0.25% trypsin-EDTA and 4mg/ml Collagenase with mechanical stress of pipetting at 30°C. Trypsin was inactivated by adding DMEM-10%FBS medium and dissociated cells were collected by centrifugation. The cells were resuspended in DMEM-10%FBS and strained by a 40um filter (Falcon). Purified single cell suspensions were subject to FACS at Rutgers Flow Cytometry Core Facility (http://rwjms1.rwjms.rutgers.edu/flow/).

### RNA-sequencing

Total RNA was immediately extracted from 100,000 EGFP-positive cells isolated by FACS using Trizol (Invitrogen). Briefly, the cell suspension was mixed with 1ml of Trizol by vigorous vortexing and kept for 5 minutes at room temperature. The mixed solution was centrifuged, and the supernatant was transferred to a new tube. Chloroform (0.2 ml) was added, vigorously vortexed, and centrifuged for 15 minutes. The supernatant was mixed with 0.5 ml of isopropanol, kept for 10 minutes at room temperature, and then centrifuged for 15 minutes. The precipitated RNA was washed by 70% ethanol and reconstituted in 30 ul water. The RNA samples were submitted to Novogene (CA, USA), converted to sequencing library, and sequenced.

### ATAC-sequencing

ATAC-sequencing was performed according to the previously described protocol^52^. Approximately 100,000 EGFP cells isolated by FACS were rinsed in PBS and resuspended in lysis buffer (10 mM Tris-HCl, pH 7.4, 10 mM NaCl, 3 mM MgCl2, 0.1% NP40). Cells were kept on ice for 10 minutes and then centrifuged at 500g for 10 minutes at 4°C. The pellets were resuspended in the reaction buffer (25 μL TD Buffer (Illumina Cat #FC-121-1030), 2.5 μL Tn5 Transposes (Illumina Cat #FC-121-1030), and 22.5 μL H_2_O). Cells were kept at 37 LJ for 30 minutes and then purified by Qiagen MinElute Kit. The DNA was eluted with 10ul of Elution buffer. Open chromatin regions flanked by the adapter sequence were amplified with 14 cycles by the thermal cycler with Illumina/Nextera i5 common adapter and i7 index adapters. The amplified sequencing library was submitted to BGI and sequencing was performed by the paired-end, 100 bp reading.

### Sequencing data analysis

The quality of RNA-sequencing data (FASTQ format) was analyzed by FastQC^70^. Then, after the adaptor sequencing primers were removed by Trimmomatic^71^, the data was mapped on zebrafish genome (GRcz10) by HISAT2^72^. The read numbers for genes were counted by HTSeq^73^ and differentially expressed genes were determined by EdgeR^74^ in R (R 4.2.2). Following differential gene expression analysis, a heatmap and volcano plot were generated using the “Heatplus” and “EnhancedVolcano” packages in R, respectively.

ATAC-Seq analysis was carried out through multiple programs in Linux. FastQC was used to get a raw quality read of the data, then low-quality reads and sequencing adapters were removed by Trimmomatic. HISAT2 mapped these reads on the zebrafish genome (GRCz10). PCR duplicates were removed with Picard (http://broadinstitute.github.io/picard/) and MACS2 was used to call peaks of open chromatin regions^75^. HOMER was used to identify Gli3 binding regions in ATAC-seq peaks and provide their gene name/specific chromosomal location^76^.

### CT scanning and quantitative analysis

To identify adult zebrafish bone phenotypes, three-month old fish (wildtype, *gli3*, and *gli;Shh* compound mutant lines) were fixed with 10% formalin overnight, stained by 0.5% phosphomolybdic acid for a week followed by water rinse several times, and micro-CT scanned at The University of Chicago (PaleoCT: http://luo-lab.uchicago.edu/paleoCT.html). The pectoral girdle bones were manually segmented, reconstructed into 3D, and visualized, using Amira 2019.4 (Fisher).

To quantify the size and shape of the pectoral girdle, twenty landmarks were used (see Extended Data Figure 8). The dorsal, anterior, and posterior extremities of the cleithrum were defined as A, G, and C, respectively. The attachment position of the scapula to the cleithrum was defined as D. The length between the dorsal and posteroventral extremities (A-D) was defined as the cleithrum height. The anterior-posterior cleithrum length was measured by extending the horizontal line anteriorly from C (B-C). The anterior-posterior coracoid length was determined by the anterior and posterior extremities of the coracoid (H-I). Following, a perpendicular line to the H-I line was drawn to maximize its length (J-K) as a coracoid height. The scapula height was determined by the dorsal and ventral extremities of the scapula (D-L). To measure the scapula width, a perpendicular line to the D-L was drawn at its center position (E-F). From the posterior view, a straight line was drawn between the dorsal and ventral edges of the glenoid fossa (M and N). Then, a perpendicular line from the M-N line to the bottom of the glenoid fossa was drawn to maximize its length (O-P; the glenoid fossa depth). The mesocoracoid height was determined by the dorsal and ventral extremities of the mesocoracoid from the posterior view (R-Q). An orthogonal line to the R-Q line at its center position was drawn as the mesocoracoid width (S-T). Each length was measured in Amira and standardized by the length of the skull height. The raw measurements are in Extended Data Table 4.

### Immunofluorescence staining

Immunofluorescence staining of EGFP and phosphorylated Smad 1/5/8 was conducted following the previously described protocol^53^. After EGFP or phosphorylated Smad 1/5/8 was stained, the plasma membrane and nucli were stained by CellMask (1/1000 diluton, Invitrogen) and DAPI (1/4000 dilution) in PBS containing 0.1% TritonX-100. EGFP antibody (Abcam #ab290) and phosphorylated Smad 1/5/8 antibody (Cell signaling #13820) were used at 1/1000 and 1/100 dilution, respectively. For the detection of the first antibody, anti-mouse Alexa 488 (Invitrogen #A21206) was used at 1/1000 dilution. The stained pectoral fin and girdles were photographed by a confocal microscope (LSM510).

### Whole-mount bone staining

Whole-mount skeletal staining of adult zebrafish with Alizarin red and Alcian blue was conducted as previously described^54^.

### Skeletal preparation of the spotted gar pectoral girdle bone

An adult, wild-caught spotted gar (*Lepisosteus oculatus*, total length was 61 cm) was euthanized with a clove oil overdose and boiled for 10 minutes in water The pectoral girdle complex was dissected out, and muscles and connective tissues were manually removed by tweezers. The cleaned pectoral girdle bones were photographed by a Canon EOS R6 camera.

### Data Deposition

Raw sequence and processed bam files of RNA-seq and ATAC-seq are available at the National Center for Biotechnology Information Sequence Read Archives (NCBI SRA), www.ncbi.nlm.nih.gov/sra (BioProject accession code no. PRJNA767802).

## Acknowledgements

We thank A. Gillis for kindly sharing elephant shark tissue. J. Lemberg provided technical assistance for µCT scanning and analysis. We thank N.H. Shubin for providing fish space, A.A. Abbasi and S. Alli for providing PXIG vector, G. Senevirathne for sharing HCR protocol, Y.J. Chang and Office of Advanced Research Computing Rutgers for computational assistance, J.J. Tena for visualization of high-throughput sequencing results, D. Remsen, S. Bennett, and staffs at the Marine Resource Center at MBL for skate embryos, J. Talbot for technical advice of live fluorescent imaging. The Bayousphere Lab (A. Ferrara) at Nicholls State University for help with gar sampling. This work was performed with Honors College with Johnson & Johnson Women in STEM2D Life Sciences Summer Undergraduate Research Fellowship (to J.W.), The Paul Robeson Scholarship, the Enrico P. Veltri Scholarship for the Life Sciences, Duncan and Nancy MacMillan Award for Research Excellence (to T.W.), the Division of Life Sciences Summer Undergraduate Research Fellowship provided by Macmillan foundation (to A.S.), The INSPIRE program (IRACDA New Jersey/New York for Science Partnerships in Research & Education) (to D.N.), NSF EDGE grant 2029216 (to I.B.), the institutional support provided by the Rutgers University School of Arts and Sciences and the Human Genetics Institute of New Jersey, the National Science Foundation under Grant IOS 2210072, Marine Biological Laboratory Whitman Fellowship (2018), Society for Developmental Biology Innovation Grant (2020), and Busch Biomedical Grant (to T.N.).

## Supplementary Information

**Extended Data Figure 1. The mutation patterns in *gli3* and *gli2b* knockout zebrafish and adult pigmentation phenotype.**

Frame shift mutations in *gli3* exon5, *gli3* exon14, and *gli2b* knockout fish, used in this paper. Green- and magenta-colored nucleotides are gRNA target sites and genetic mutations, respectively.

**Extended Data Figure 2. The survival ratio of wildtype and *gli3*^14ins/14ins^ juveniles.**

One clutch of fertilized eggs from a mating pair of wildtype fish and two clutches of fertilized eggs from two mating pairs of *gli3*^14ins/14ins^ fish were obtained. They were raised up until 49 days and the number of living juveniles was counted at each time point. The mortality of wildtype and *gli3*^14ins/14ins^ juveniles did not show significant difference.

**Extended Data Figure 3. HCR of *sp7* and *col2a1a* in wildtype and BMS-833923 treated embryos.**

**a** and **b**) confocal-microscopy imaging of embryos stained by HCR with *sp7* (osteoblasts in the cleithrum) and *col2a1a* (chondrocytes in the scapulocoracoid) probes in wildtype (**a**) and BMS-833923-treated embryos (**b**) at 72 hpf. Images are maximum intensity projections of the lateral views of the embryos. In BMS-833923 treated embryos, the *sp7* staining is weaker in the dorsal extremity and middle portion of the cleithrum (arrows) than those of wildtype embryos. **c**) The number of chondrocytes in the scapulocoracoid of *gli3*^+/+^, *gli3*^14ins/14ins^, and BMS-833923 treated embryos. The number of chondrocytes increases in *gli3*^14ins/14ins^ and BMS-833923 embryos (*N*= 19, 18, and 18 for *gli3*^+/+^, *gli3*^14ins/14ins^, and BMS-833923 embryos, respectively). The scale bar is 300 μm. ** and * indicate *p*<0.01 and *p*<0.05 in one tailed t-test, respectively. *p*= 0.0011 between *gli3*^+/+^ and *gli3*^14ins/14ins^ embryos and *p*=0.0329 between *gli3*^+/+^ and BMS-833923 treated embryos.

**Extended Data Figure 4. The expression pattern of *activin receptor type 1* in the pectoral fin of skates (*L.erinacea*).**

**a** and **b**) *Acvr1* is expressed in the pectoral fin and the body trunk at embryonic stage 30 (**a**; ISH with sense probe and **b**; with antisense probe). The scale bar length is 1 mm.

**Extended Data Figure 5. Bone staining of *gli3*^14ins/14ins^ fish and pectoral girdle phenotype.**

**a-c, g**, and **h**) Alcian blue and alizarin red staining of wildtype (**a**,**c**, and **g**) and *gli3*^14ins/14ins^ fish (**b**, **d**, and **h**). In *gli3*^14ins/14ins^, ectopic ossification is observed in the eye lens (5/6 individuals), indicated by white arrows in **b**). The skull roof bones of *gli3*^14ins/14ins^ fish are more mineralized than those of wildtype fish (**a** and **b**). The number of pectoral fin rays do not show significant difference between wildtype and *gli3*^14ins/14ins^ fish (**c**, **d**). The pectoral fin on one random side of the body shifts dorsally due to the cleithrum defect in severely affected *gli3*^14ins/14ins^ fish (**e**, **f**, **g**, **h**). On one side, the pectoral fin position is normal (**e**, black arrow), and the fin on the other side attaches dorsally to the body (**f**, white arrow). All scale bars are 1.25 mm.

**Extended Data Figure 6. Quantitative analysis of the pectoral girdle by 3D reconstruction.**

Red dots indicate landmarks used to quantify pectoral girdle bone morphology. The method to set the landmarks and quantify the line length that connects them is described in Method. **a**) the lateral view of the wildtype left pectoral girdle. **b**) the posterior view of the wildtype left pectoral girdle bones. A; the dorsal extremity of the cleithrum. B and C; the anterior and posterior ends of the dorsal shaft of the cleithrum. D; the attachment of the scapula to the cleithrum. E and F; the anterior and posterior ends of the scapula. G; the anterior extremity of the cleithrum. H; the anterior extremity of the coracoid. I; the posterior extremity of the coracoid, J and K; the dorsal and ventral ends of the widest point of the cleithrum. L; the ventral extremity of the scapula. In this paper, the following distance were measured to compare pectoral girdle morphology among different genotypes. A-D; the cleithrum height, B-C; the cleithrum width, G-D; the cleithrum length, D-L; the scapula length, E-F; the scapula width, H-I; the coracoid length, J-K; the coracoid width. M, N; the dorsal and ventral edges of the glenoid fossa. O, P; the ends of the line orthogonal to the M-N line. R, Q; the dorsal and ventral extremities of the mesocoracoid. S,T; the left and right ends of the line orthogonal to the R-Q line. All measurement is summarized in Extended Data Table 3.

**Extended Data Figure 7. The modest phenotype of the cleithrum in *gli3*^14ins/14ins^ fish.**

While the pectoral fin on one side in approximately 20 % of *gli3*^14ins/14ins^ fish shows the dorsal shift, the pectoral fins on the other side of these fish and pectoral fins on both sides in the remaining 80% fish show the modest cleithrum phenotype. The modestly affected cleithrum in *gli3*^14ins/14ins^ fish is dorsoventrally shorter (black arrows) and anteroposteriorly wider (white arrows) than that of wildtype fish.

**Extended Data Figure 8. Skeletal preparation of adult spotted gar (*Lepisosteus oculatus*) pectoral girdle.**

The lateral view (**a**) and medial view (**b**) of the pectoral girdle. cl; cleithrum, clmw; medial wing of cleithrum, r; fin rays, s; scapula, scl; supracleithrum, scldp; dorsoanterior process of supracleithrum, sclvp; ventroanterior process of supracleithrum. Note that the supracleithrum is detached from the cleithrum. Scale bar = 1 cm.

**Extended Data Table 1. PCR primers used to create gRNAs, conduct T7 assay, and clone genes.**

**Extended Data Table 2. Upregulated and Downregulated gene lists identified by RNA-seq in *gli3*^14ins/14ins^ pectoral fin.**

**Extended Data Table 3. The list of wildtype ATAC-seq peaks annotated to genes.**

ACRs called by MACS^79^ were annotated to genes depending on the proximity to transcription start sites (see Method).

**Extended Data Table 4. Quantification data of pectoral girdle morphology.**

The distance between landmarks of 3D-reconstructed pectoral girdle bones in wildtype, *gli3*^14ins/14ins^, and *gli3*^14ins/14ins^; *shha*^30%^ fish were measured in Amira. The landmark points for each measurement were defined in Extended Data Figure 8.

## Author Contributions

T.W., I.B, and T.N. designed research. K.F. and T.N. created genetically modified fish. J.W., A.S., and T.N. analyzed RNA-sequencing data, ATAC-sequencing data, and conducted chromogenic whole mount *in situ* hybridization and HCR. A.E., A.A. and K.F genotyped transgenic and knockout fish. T.S. and T.N. conducted CT scan. T.W. created pectoral girdle bone 3D-reconstructions from CT scan data. D.N. conducted in situ hybridization of skate embryos. T.W., J.W., and T.N. analyzed the data. S.K. and T.N. wrote the paper.

