## Supplemental Table 1 for "Distinct ossification trade-offs illuminate the shoulder girdle reconfiguration at the water-to-land transition"

CRISPR gRNA oligos

zebrafish *gli3*_gRNA_exon5_F

5'- AATTAATACGACTCACTATAGGTACTGGATGATGAAAAGCGTTTTAGAGCTAGAAATAGC -3'

zebrafish *gli2b*_gRNA_F

5'- AATTAATACGACTCACTATAGGTTCCGATGGCCGGCCATCGTTTTAGAGCTAGAAATAGC -3'

common_gRNA_R

5'- AAAAGCACCGACTCGGTGCCACTTTTTCAAGTTGATAACGGACTAGCCTTATT TTAACT TGCTATTTCTAGCTCTAAAAC -3'

T7 assay PCR primers

zebrafish *gli3*_exon5_cont_F

5'- TGTCTTTCTGTCAGGTTGTAAAAA -3'

zebrafish *gli3*_exon5_cont_R

5'- CTCCTTTACCCGTCGGTTTT -3'

PCR product: 527 bp

zebrafish *gli2b*_cont_F

5'- CAACAACAAAATAATTCAAGCACT-3'

zebrafish *gli2b*_cont_R

5'- CTGCCTGCCTGTTTAAAGACT-3'

PCR product: 804 bp

Genotyping primers

zebrafish *gli3*_exon5_genotyping_F

5'- CTGCAGAAACCATGGACAAA -3'

zebrafish *gli3*_exon5_genotyping_R

5'- CCTTCAATGGGGACAGACTC -3'

PCR product: 112 bp

zebrafish *gli2b*_genotyping_F

5'- GACCGCAACGGATCAATTA-3'

zebrafish *gli2b*_genotyping_R

5'- GTGAAGCTTCTGGCCAGTGT-3'

PCR product: 932 bp

Gene and enhancer cloning primers

Elephantshark_*CNE14*_F

5'-GCTTACGACACCTACCAATG -3'

Elephantshark_*CNE14*_R

5'- CACCATCTCCGGAACTACAC -3'

Zebrafish *acvr1l*_F

5'--3'

Zebrafish acvr1l_R

5'- TCACGTCTTCCTGCA -3'

Skate_*acvr1*_F

5'-TGCTGGAAGAGCACATTTTG -3'

Skate *acvr1*_R

5'- ATGGGCCACTACTGTCCTTG-3'

Extended Data Table 1.
