## Supplementary figures for "Distinct ossification trade-offs illuminate the shoulder girdle reconfiguration at the water-to-land transition"

##### *gli3* exon 5 $^{14ins/14ins}$

WT; TTCACATCCTTACATTAACCCC - - - - - - - - - - - - - - TACATGGACTACATACGCTCCTTGCA

MT; TTCACATCCTTACATTAACCCCTACATACTACATACTACATACGCTACATACGCTCCTTGCA

##### *gli2b* $^{17ins/17ins}$

WT; GTCTCCAGCGGATTATTACCACCTG - - - - - - AT - - - - - - - - GG - - -CCGGCCATCGGAACC

MT; GTCTCCAGCGGATTATTACCACCTGGGGATTATTACCATACGGGTTCCGGCCATCGGAACC

Extended Data Figure 1

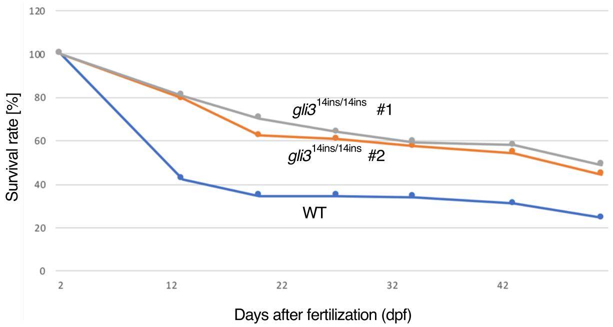


### Extended Data Figure 2

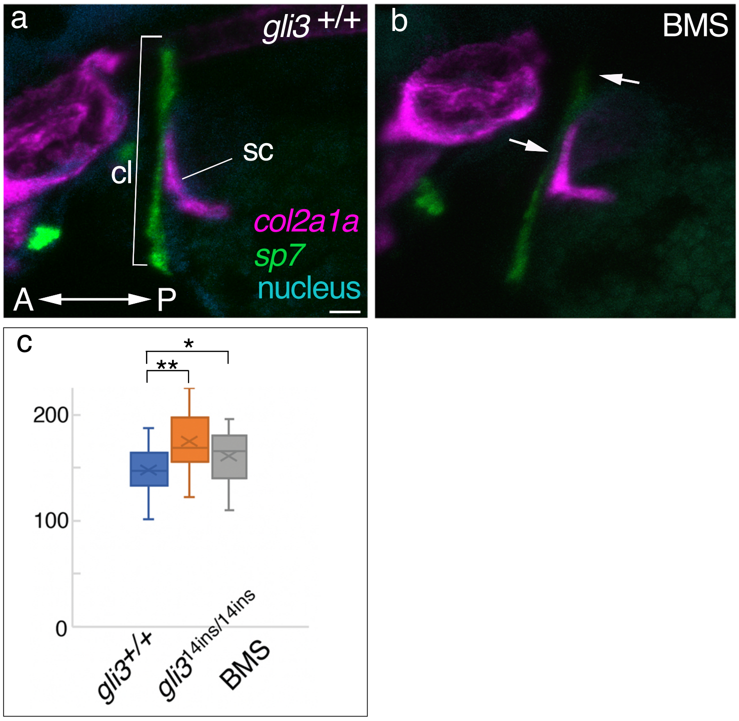


Extended Data Figure 3

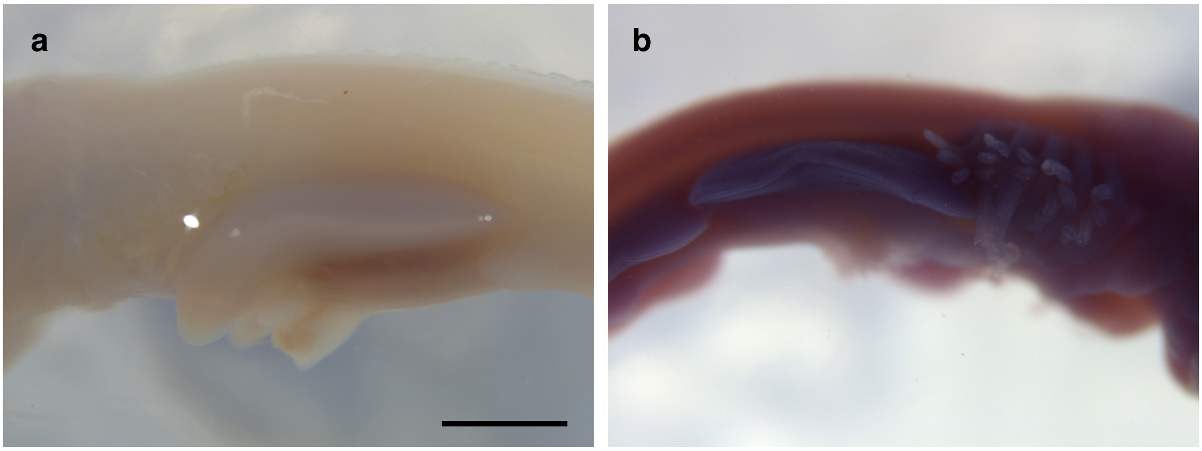


**Extended Data Figure. 4**

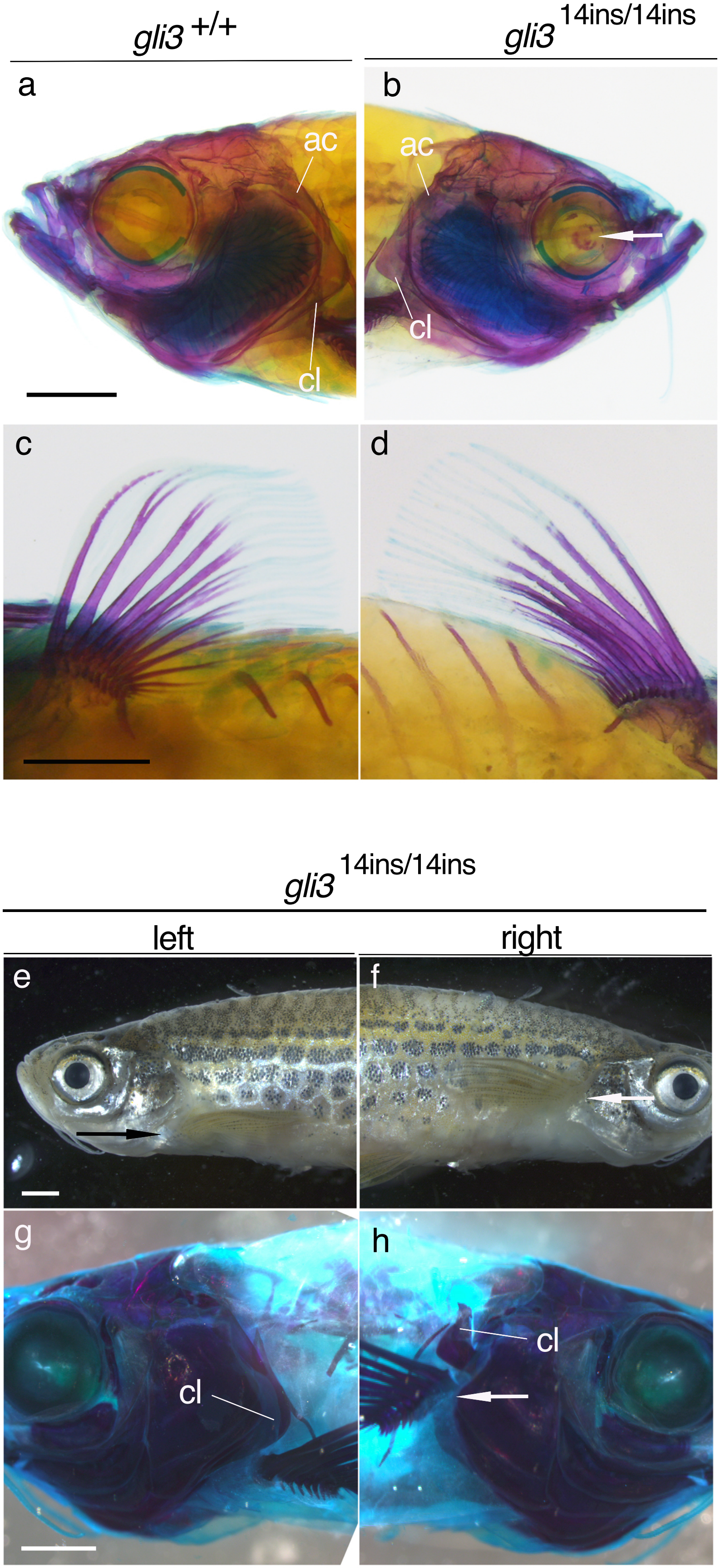


Extended Data Figure 5

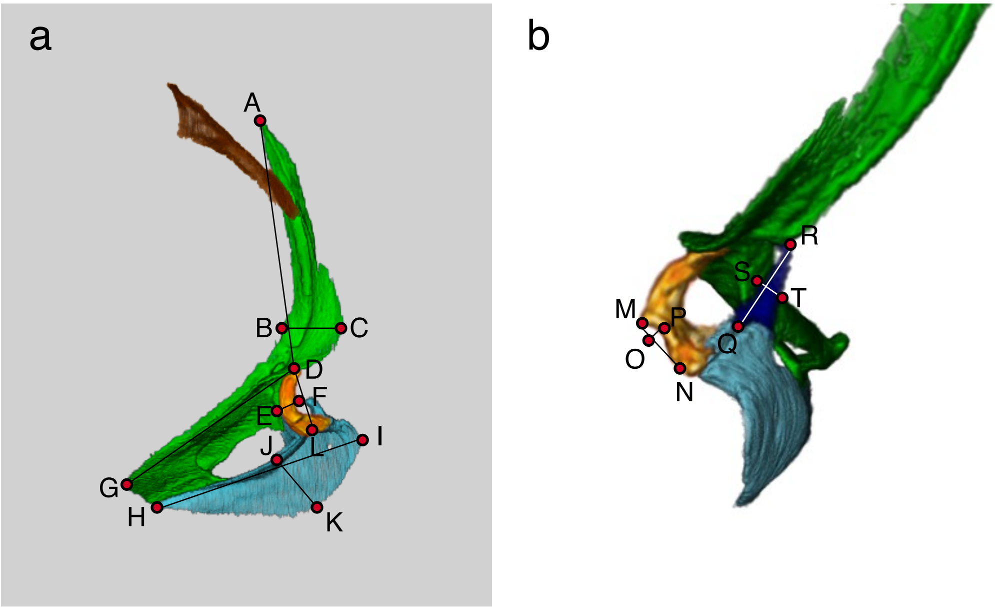


Extended Data Figure 6

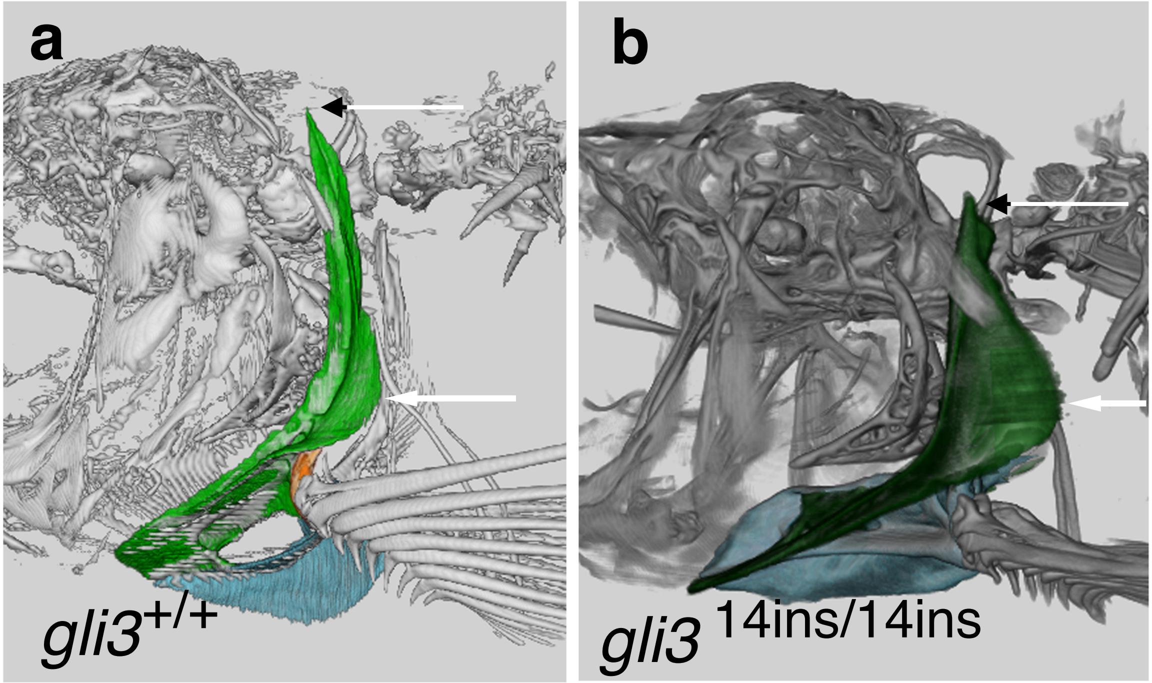


Extended Data Figure 7

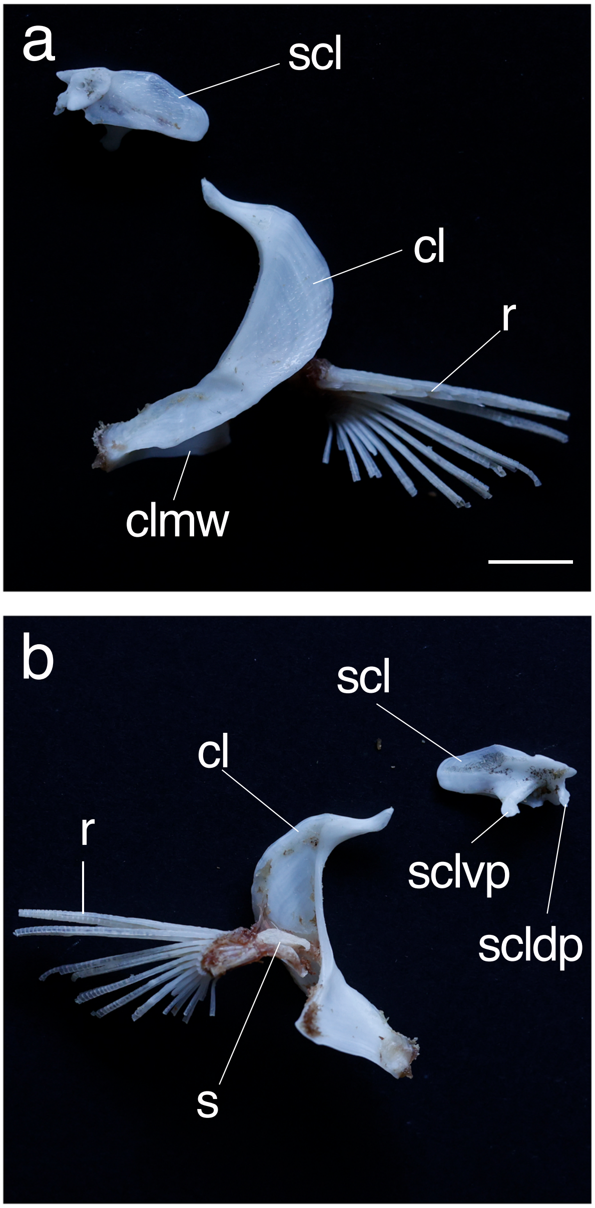


Extended Data Figure 8
